# A Spectroscopic Approach to Unravel the Local Conformations of G-quadruplex Using CD-active Fluorescent Base Analogues

**DOI:** 10.1101/2022.07.06.499065

**Authors:** Davis Jose, Miya Mary Michael, Christopher Bentsen, Brandon Rosenblum, Adriana Zelaya

## Abstract

The formation of a stable G-quadruplex (GQ) can inhibit the elevated telomerase activity that is common in most cancers. The global structure and the thermal stability of the GQs are usually evaluated by spectroscopic methods and thermal denaturation properties. However, most biochemical processes involving GQs might require local conformational changes at the guanine tetrad (G4) level. These local conformational changes of individual G4 layers during protein and drug interactions have not yet been explored in detail. In this study we monitored the local conformations of individual G4 layers in GQs using 6-methylisoxanthopterine (6MI) chromophores, which are circular dichroism (CD)-active fluorescent base analogues of guanine, as local conformational probes. A synthetic, tetra-molecular, parallel GQ with site-specifically positioned 6MI monomers or dimers was used as the experimental construct. Analytical ultracentrifugation studies and gel electrophoretic studies showed that properly positioned 6MI monomers and dimers could form stable GQs with CD-active fluorescent G4 layers. The local conformation of individual fluorescent G4 layers in the GQ structure was then tracked by monitoring the absorbance, fluorescence intensity, thermal melting, fluorescence quenching and CD changes of the incorporated probes. Overall, these studies showed that site-specifically incorporated fluorescent base analogues could be used as probes to monitor the local conformational changes of individual G4 layers of a GQ structure. This method can be applied to explore the details of small molecule-GQ interaction at the level of the individual G4 layers, which may prove useful in designing drugs to treat GQ-related genetic disorders, cancer and aging.

## INTRODUCTION

G-quadruplexes (GQs) are stable, noncanonical nucleic acid secondary structures that are found mainly at telomeres and other guanine-rich sequences, such as promoter regions of DNA and untranslated regions of RNA[1–5]. Telomeres, the multiple repeats of guanine-rich sequences, serve as protective structures at the ends of linear chromosomes and play a significant role in stabilizing the genome[6]. Cellular replication causes telomere shortening, which leads to replicative senescence and apoptosis. However, in most cancer cells this natural process is eluded by upregulating the reverse transcriptase telomerase that elongates telomeres [7]. Effective telomere elongation requires unfolding of GQ secondary structures by specific helicase enzymes. The stabilization of GQ structures by various methods can inhibit the unfolding of G-quadruplex structures and results in preventing extensive telomere elongation.

GQs may form in single-stranded DNA and RNA G-rich sequences under physiological conditions. Hoogsteen-type base-pairing forms a square planar guanine tetrad (Figure 1 C), and the stacking of two or more of these tetrads leads to GQ formation. Monovalent cations of a specific size range occupy the central cavity of the G4 layers within GQs, relieving the repulsion between oxygen atoms and thus playing a significant role in the formation and stability of GQs [8, 9]. Thermal denaturation assays at different salt concentrations are typically used to characterize the stability of GQs formed from various single-stranded (ss) DNA sequences[10–13].

**Figure 1:**
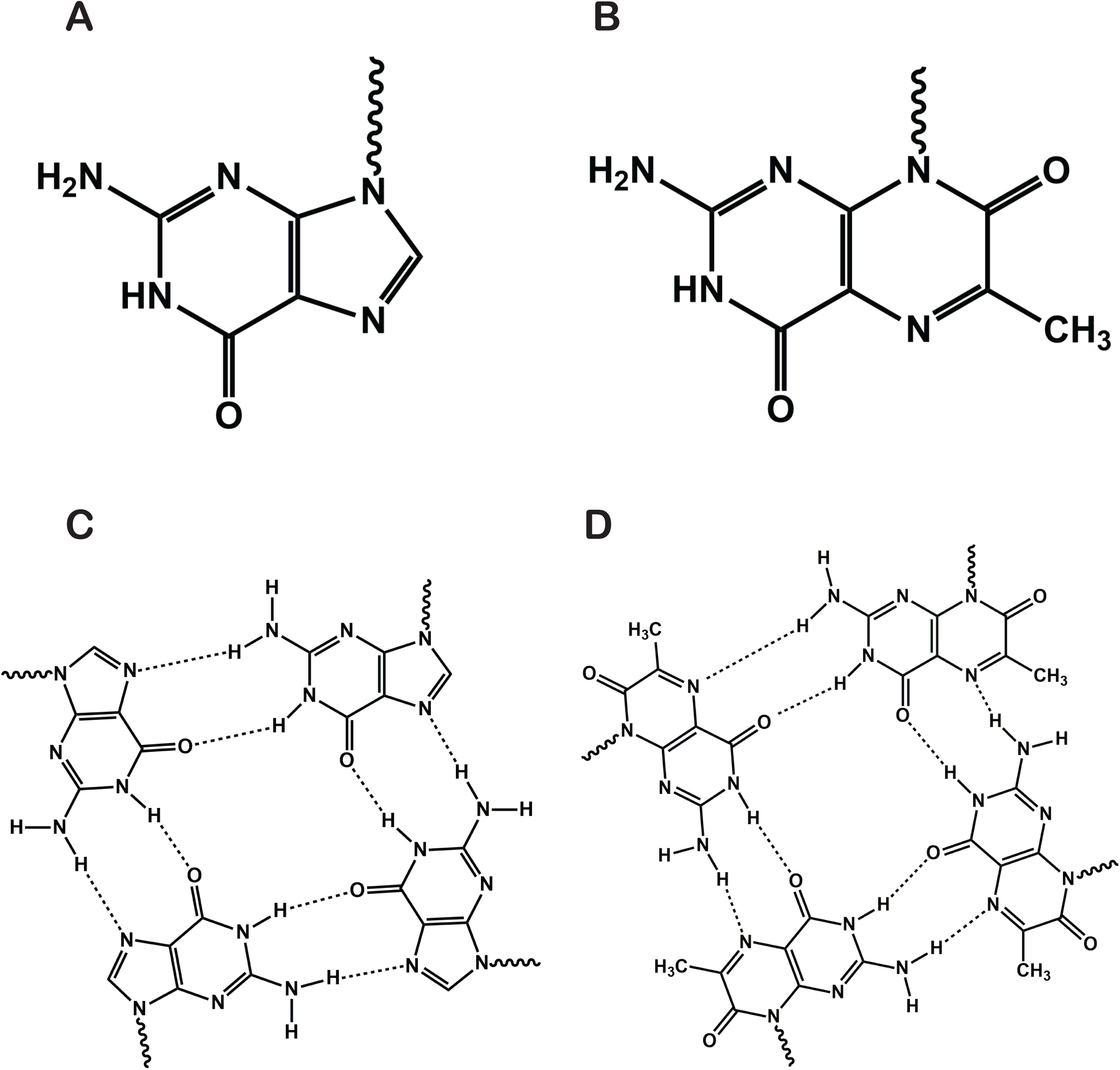
Chemical structures of the bases and the corresponding G4 layers. (A) Guanine, (B) 6MI, (C) Structure of the G4 layer formed by Guanines (D) Proposed structure of the G4 layer formed by 6MI. Potential hydrogen bonding interactions are shown in dashed lines.

ssDNA sequences with the potential to form GQs are not limited to the guanine-rich telomeric region, but are also abundant in other guanine-rich regions of the genomes. Within the human genome, computational studies have identified more than 700,000 potential quadruplex-forming sequences (PQS) clustered at specific functionally active genomic loci, including telomeres, oncogene promoters, immunoglobulin switch regions, DNA replication origins and recombination sites[14]. In RNA, GQs have been found in mRNAs and non-coding RNAs, indicating the potential of RNA GQs to regulate pre- and post-transcriptional gene expression[4]. The high stability, structural versatility, and functional diversity of GQs have enabled their use in many medical and biotechnology applications. Apart from being targets for anti-cancer anti-aging drugs[15–18], GQs are used as biosensors and molecular switches[19, 20].

Even though global circular dichroism (CD) and fluorescence intensity changes have been previously used to track the conformational changes in GQs[21–23], the UV absorbance and CD signals of individual G4 layers are masked by spectral contributions from the canonical nucleotide residues of the DNA. As a consequence, the details of the stabilizing interactions at the individual G4 level of the GQs are not yet well defined. In this study we have utilized three different, but complementary, spectroscopic methods involving the CD-active fluorescent guanine analogue, 6MI, which have been position-specifically placed into the nucleic acid framework to study local conformations and stabilizing interactions at the G4 level within GQ constructs containing a number of G4 components.

6MI is a fluorescent base analog (Figure 1B) of the canonical base guanine (Figure 1A) that can be incorporated into nucleic acid constructs as either a monomer or a dimer conformational probe to study the local environments of specific guanine residues in DNAs and RNAs. It was previously shown that similar to the commonly used fluorescent adenine analog 2-Aminopurine (2AP), incorporation of a single 6MI residue into a double-stranded (ds) DNA has little effect on the thermal stability of the DNA duplex[24, 25]. The 6-MI nucleotides exhibit an excitation maximum at ∼ 350 nm and an emission peak around ∼450 nm. Once incorporated into a DNA sequence, the fluorescence emission maximum shifts to ∼ 430 nm, and fluorescence quenching is observed. The fluorescence intensity is further quenched in duplex DNA sequences, and this has been attributed to the stacking interactions of the probe with neighboring bases. In general, it was found that maximum quenching was observed with guanine as the nearest neighbor, whereas the least fluorescent quenching was observed when thymine residues served as neighboring bases, although some exceptions to this generalization have been observed [26]. The fact that the fluorescence signal of 6-MI is sensitive to base-stacking and nucleic acid sequence context prompted us to use this probe to monitor the local conformations of individual G-tetrad layers in a GQ structure. The hypothetical structures of G4 layers containing guanines and 6MI residues instead of guanines are shown in Figure 1C and D respectively. The proposed hydrogen bonding pattern suggests that under favorable conditions, four 6MI bases can form a stable G4 layer similar to that formed with guanines.

The uniqueness of the position-specific CD-active fluorescent base analog approach in studying the conformation of nucleic acids is that it allows the simultaneous tracking of: (i) the global conformational changes of the nucleic acid framework; and (ii) the more unique local conformational changes associated with individual bases at defined locations within the GQ, without optical interference from canonical nucleic acid bases or protein residues, because the latter are effectively ‘transparent’ at the wavelengths used in this study[27–29]. This approach was successfully employed in studies where position-specific DNA breathing properties of DNA duplex and fork constructs were tracked. Further, this method was used in defining the local changes in the conformations of ssDNA, P/T junctions, fork junctions and duplex regions as a function of probe position in biological processes such as DNA replication, transcription and specific and nonspecific protein binding to target sites on nucleic acid constructs [30–33].

There are studies in the literature where the fluorescence properties of CD-active fluorescent base analogues, mainly 2-Aminopurine (2AP), a fluorescent base analogue of adenine, have been used to track the formation and stability of GQs [19,34–40]. In these studies, the fluorescent probes were either positioned at a secondary structure loop or used as a single substitution in specific G4 layers, but only monitored at the level of global circular dichroism and fluorescence intensity changes. The main drawback of these studies was the lack of information at the level of individual G4 layers of GQs.

In the present study, in order to better understand the formation, stabilities and interactions of individual G4 layers, we incorporated one or more CD-active fluorescent G4 layers directly into the GQ structure. Our results show that site-specifically positioned 6MIs can form a stable GQ containing one or more fluorescent G4 layers. Furthermore, we showed that this approach can be used for the simultaneous determination of global and local conformations and stabilities of GQs and the individual G4 layers they contain. This approach will help us to analyze and understand the interactions of GQ with GQ-related proteins and therapeutically significant small molecules, which should eventually lead to the design of better drugs for GQ-related diseases.

## MATERIALS AND METHODS

### DNA constructs, nomenclature, and annealing

Unlabeled DNA oligonucleotides were purchased either from Integrated DNA Technologies (IDT) (Coralville, IA, USA) or from Fidelity Systems (Gaithersburg, MD, USA). 6MI labeled oligonucleotides were purchased from Fidelity Systems (Gaithersburg, MD, USA). All oligos had -OH groups at both the 3’ and the 5’ ends of the ssDNA chains. Lyophilized DNA was re-suspended in deionized water and concentrations were determined by UV absorbance at 260 nm (25°C), using extinction coefficients furnished by the manufacturer. The sequences and nomenclature of the DNA constructs used in this study are listed in Table 1. Tetramolecular GQs were formed by heating 50 μM of the ssDNA in 10mM Tris, 100mM (or 50mM) KCl, pH 7.5 at 95°C for 15 minutes, followed by overnight cooling at room temperature.

**Table 1.** Nomenclature, Sequence and molar mass of the of 6MI containing ssDNA constructs and the expected molar mass of the GQ formed from each construct. (1Da = 1g/mol)

| Construct | Sequence | Molar mass of oligomer | Expected Molar mass of GQ |
| --- | --- | --- | --- |
| 8G | 5'-TTAGGGGGGGGA<sub>15-3'</sub> | $8.19 \times 10^4$ g/mol | $3.28 \times 10^4$ g/mol |
| 8Gm | 5'-TTAGGGG6MIGGGA<sub>15-3'</sub> | $8.23 \times 10^4$ g/mol | $3.29 \times 10^4$ g/mol |
| 8Gd | 5'-TTAGGGG6MI6MIGGA<sub>15-3'</sub> | $8.28 \times 10^4$ g/mol | $3.31 \times 10^4$ g/mol |

**Table 2.** T_m_ Values for GQ constructs from Fluorescence and UV thermal denaturation curves. Summary table showing the melting temperatures of GQs formed from 8Gm construct at different salt concentrations and their respective error (standard deviation). The data are extracted from Figures 5 and 7.

| Construct | Salt concentration | T<sub>m</sub>, °C ($\Delta A_{350nm}$) | T<sub>m</sub>, °C ($\Delta F_{430nm}$) |
| --- | --- | --- | --- |
| 8Gm | 10mM KCl | Not detectable | $47.5 \pm 1$ |
| 8Gm | 50mM KCl | $47.5 \pm 1$ | $48 \pm 1$ |
| 8Gm | 100mM KCl | $50.5 \pm 1$ | $49.5 \pm 1$ |

**Table 3.**
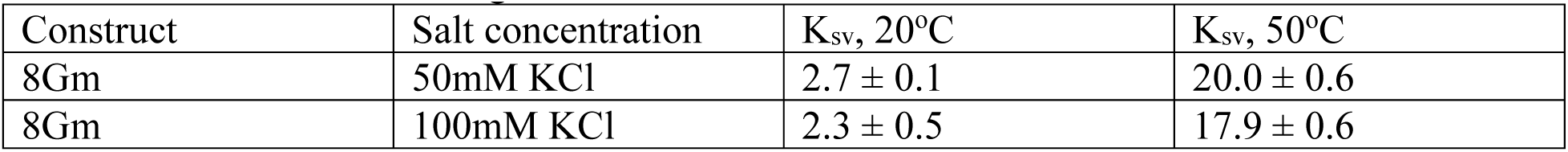
Stern-Volmer titration fit parameters for the Acrylamide fluorescence quenching experiments. Summary table showing the Stern-Volmer constants of GQs from 8Gm constructs at different salt concentrations and temperatures with their respective error (standard deviation). The data are extracted from Figure 8.

### Spectroscopic measurements

UV-visible spectra were obtained with a Varian Cary 3E UV-Visible spectrophotometer equipped with a Peltier temperature controller using samples in a 1 cm path-length quartz cuvettes and a DNA oligomer concentration of 5 μM. For UV-visible thermal melting measurements, the absorbance of the samples was tracked at 260 nm, which is near the absorbance maximum of the canonical bases and, at 350nm, near the absorbance maximum of 6MI. CD spectra were measured in 1 cm path-length optical grade quartz cells over the wavelength range 230-500 nm using a JASCO model J-1500 CD spectrophotometer equipped with a Peltier temperature controller. For each spectrum, 3 to 5 individual data sets were collected at a bandwidth of 0.5 nm and a scanning rate of either 20 nm/min or 50 nm/min. The experiments were performed at a DNA lattice concentration of 2 μM. Fluorescence measurements were made using 4×4 mm or 2×2 mm optical quartz cells in an Edinburgh FS5 spectrofluorometer. For the fluorescence intensity measurements, samples were excited at 350 nm, and emission spectra were collected from 360 nm to 500 nm. For the fluorescence melting measurements, the temperature of the holder was tracked and the samples were excited at 350 nm, and the emission peak was monitored near the emission maximum of 430 nm. For fluorescence quenching experiments, samples containing DNA constructs were titrated with increasing amounts of acrylamide (1M) stock solution. In all experiments the samples were gently mixed, equilibrated for two minutes, and scanned at either 20°C or 50°C. Corrections for background fluorescence were made by subtracting the fluorescence intensities obtained with an unlabeled ssDNA construct of the same length.

### Data Analysis

The absorbance spectra were baseline corrected with the buffer and plotted as absorbance as a function of wavelength. The CD spectra replicates were averaged, buffer-subtracted and plotted as graphs of Δε (M^−1^cm^−1^) as a function of wavelength, where molarity refers to that of the oligomer construct. Raw fluorescence data were corrected for background counts and dilution and spectral contributions from unlabeled ssDNA. Fluorescence normalization was done using the following equation;

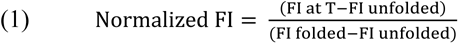

Here FI unfolded is the average value of fluorescence at the highest 5°C and FI unfolded is the average value of fluorescence at the lowest 5°C. All melting curves and the accompanying statistics were generated using OriginPro plotting and melting temperatures were determined manually and by the first derivative method, and both values were always confirmed by cross checking. Error bars for each data point represent the standard deviation of three to four repeats of the relevant measurement.

Acrylamide fluorescence quenching data were plotted as ratios of the fluorescence intensity in the absence of a quencher (*F*_0_) over the intensity (*F*) after the addition of quencher (*Q*) [41]. The data points showed a clear linear trend and were fit by linear regression to obtain *K_SV_*, the Stern-Volmer quenching constant, using the Stern-Volmer equation written as follows:

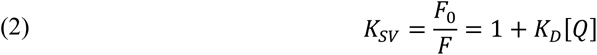

Here *K_D_* is the Stern-Volmer quenching constant (for collisional quenching) and [Q] is the quencher concentration (M) (here monomeric acrylamide). We note that *K_D_^-1^* is the quencher concentration at which 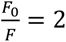 (i.e., the concentration at which 50% of the initial fluorescence has been quenched. All the graphs and the accompanying statistics were generated using OriginPro plotting and fitting software. Error bars for each data point represent the standard deviation of three to six repeats of the relevant measurement.

### Analytical Ultracentrifugation and Electrophoresis Mobility Shift Assay

Sedimentation velocity experiments were performed in a Beckman Optima XL-I Analytical Ultracentrifuge with a Beckman An60Ti ultracentrifuge rotor. Unless otherwise stated, experiments were run at 20°C at a rotor speed of 50,000 rpm and monitored by UV absorbance at 260 nm. In each experiment, 400 μl of sample and 410 μl of reference buffer were loaded, respectively, into the sample and reference sectors of a 1.2-cm double-sector Epon centerpiece in a standard analytical ultracentrifuge cell. Sedimentation velocity data were collected continuously over periods of up to 8 h. All data analyses for the velocity sedimentation experiments were performed using the SedFit program [42–44]. The root-mean squared-deviations (rmsd) for all sedimentation velocity experiments were 0.008 or less. Sedimentation results were plotted as molar mass distribution c(M) plots of the relative concentrations of each size component versus molecular weight (Da).

For the Electrophoresis Mobility Shift Assay (EMSA), the ssDNA oligonucleotides were 5’ end-labeled with [γ-^32^P] ATP, and tetramolecular GQs were formed by heating in 10mM Tris, 100mM (or 50mM) KCl, pH 7.5 at 95°C for 15 minutes followed by overnight cooling at room temperature. The folded DNA constructs were added to the native gel loading buffer (0.25% bromophenol blue, 0.25% xylene cyanol, and 30% glycerol) and electrophoresed on a nondenaturing 12% polyacrylamide gel for 3 to 5 hours at 20mA at room temperature. Both buffer and gel contained the same constituents as those in which the DNA was folded. The gel was dried and exposed to a PhosphorImager screen (Molecular Dynamics) for 8 to 14 hours.

## RESULTS AND DISCUSSIONS

### Constructing a fluorescent G4 layer with guanine analogue 6MI

Probing the local conformational details of nucleic acid lattices is critical for developing an understanding of phosphate backbone orientations and stacking perturbations of nucleotides in various nucleic acid sequences and conformations. To monitor the formation, stability, and local conformation of individual G4 layers, we incorporated either a monomer or a dimer probe of 6MI, a fluorescent base analogue of guanine, into a synthetic ssDNA sequence that is known to form stable GQs [45]. The chemical structures of guanine, 6MI and the G4 layers formed within GQ constructs using possible Hoogsteen base pairing are shown in Figure 1, and the sequences used in our studies are summarized in Table 1. The synthetic ssDNA strand denoted as 8G is known to form a very stable parallel GQ with eight layers of G4s [45]. In Figure 2 we shown the structures of GQ formed by 8G constructs and by the modified constructs 8Gm and 8Gd where the G residues at the 5^th^ or 5^th^ and 6^th^ positions are replaced by the 6MI fluorescent analogue probe. We hypothesize, since 6MI tetrads can form similar Hoogsteen-base pairing patterns as guanines, that the 6MI-labeled ss DNA constructs, under GQ forming conditions, can form a stable GQ with either one or two CD-active fluorescent G4 layers (Fig 2B, C). Based on this assumption, monitoring the absorbance, fluorescence, and circular dichroism of these CD active-fluorescent G4 layers should reveal additional aspects of the structural and molecular features of GQs that have been previously unexplored.

**Figure 2.**
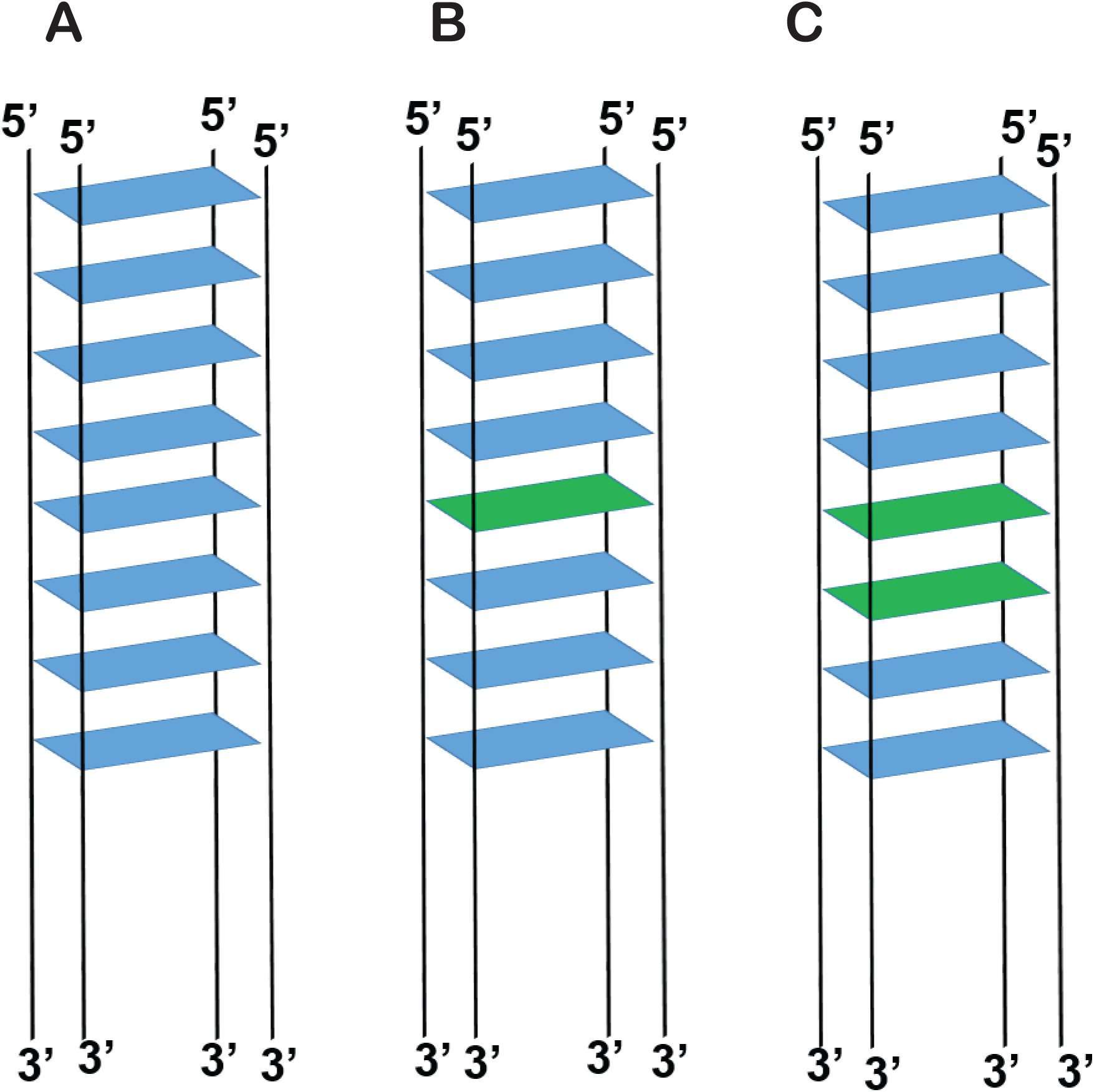
Scheme of G-quadruplexes with or without site specifically positioned fluorescent probes. In panels A, B, and C, Guanine tetrads are shown in blue and 6MI tetrads in Green. GQ formed by (A) 8G (5’-TTAG_8_A_15_-3’), (B) 8Gm (5’-TTAG_4_6MIG_3_A_15_-3’), where 6MI monomer is inserted at position 5 from the 5’ end (C) 8Gd (5’-TTAG_4_6MI6MIG_2_A_15_-3’) where 6MI dimer is inserted at positions 5 and 6 from the 5’ end.

### Electrophoresis mobility shift assays (EMSA) and sedimentation velocity experiments show that site-specific incorporation of the fluorescent guanine analogue 6MI containing ssDNA strands can form a stable GQ structure

Electrophoretic mobility shift assays were performed on unlabeled (8G), 6MI monomer labeled (8Gm) and 6MI dimer labeled (8Gd) constructs to determine whether 6MI incorporated DNA constructs can form stable GQ constructs (Figure 3A). A T_25_mer was used as a control to monitor the approximate gel-shift position of residual unhybridized ssDNA strands. The unlabeled ssDNA construct (8G) showed three distinct bands, whereas the 8Gm and 8Gd constructs showed only two bands. The bottom band that matches the T_25_ control was seen for all three components and presumably corresponds to the unhybridized ssDNA strands that did not take part in the formation of GQ.

**Figure 3.**
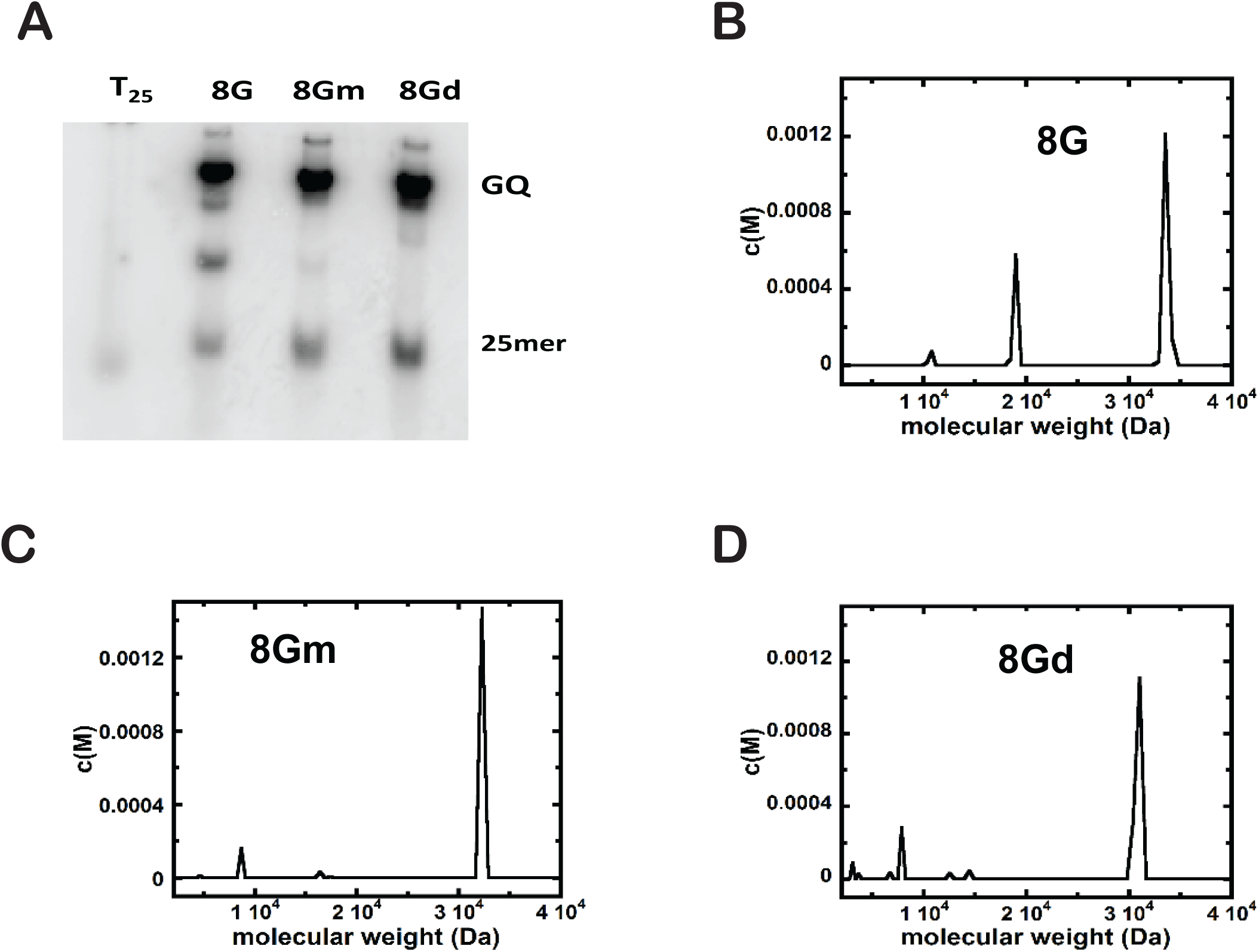
Conformation of GQ formation by 6MI incorporated strands using gel electrophoresis and sedimentation velocity experiments. (A) Top band in electrophoresis shows the formation of GQ with all constructs. The peak at ∼ 3.2 KDa in sedimentation velocity experiments confirmed the formation of GQ with (B) 8G, (C) 8Gm, (D) 8Gd. All experiments were performed in 10mM Tris, 100 mM KCl, pH 7.5.

The middle band, seen only in the unlabeled 8G construct, is attributed to an intermediate, most probably a bimolecular duplex structure which might be formed due to the extensive G-G Hoogsteen base pairing, that might serve as a precursor in the formation of the tetramolecular QC structure. This middle band is not observed with the 6MI monomer- (8Gm) or dimer-(8Gd) labeled constructs, suggesting incorporation of 6MI prevents formation of partially stable duplex structures and only the final tetramolecular GQ products are formed. The absence of this intermediate structure suggests that the insertion of the 6MI probe(s) might favor the formation of GQ. One possible interpretation for this observation is that the disruptions in the G-G run by the incorporation of 6MI probes prevents the formation of partially stable duplex structure. Another possibility is that the larger surface area of 6MI might favor more efficient base stacking due to the increased π-π interaction, thus enhancing the formation of stable GQs, meaning that the stacking of 6MI residues with guanine or with other 6MI residues forms more stable intermediates in QC structure formation than does guanine-guanine stacking, thus driving the conversion of all the intermediately formed bimolecular constructs into tetramolecular GQ structures. A similar observation was made by Gros et.al, in that this group found quadruplex formation appears to proceed more rapidly with ssDNA strands containing 6MI near the 5’end of the parallel strand than with the unlabeled ssDNA strands [39]. The top band was present in all constructs, and it was assigned to the fully formed GQ structures. However, in the major top gel band, we notice that there is a very small, but detectable, additional band just beneath the main band in all cases. We attribute this small band to a very low concentration of trimer, which may be responsible for this ‘spreading’ of the major tetramer band.

Analytical ultracentrifugation was used to further characterize and confirm the formation of the final tetrameric GQ structures from the component ssDNA strands. The molecular mass of all the oligomers and the expected molecular mass of the corresponding GQs formed from them are shown in Table 1. Sedimentation velocity profiles of all constructs were obtained at the input molarity of the individual ssDNA concentrations of 3 μM. A small peak that corresponds to a molecular weight of ∼ 1.0 x10^4^ Da is observed in all experiments, which is attributed to residual unhybridized ssDNA constructs. The unlabeled DNA construct (8G) showed two significant peaks with molecular weights of ∼ 1.9 x10^4^ Da and ∼ 3.4 x10^4^ Da, respectively (Figure 3B). The observed molecular weight for the first significant peak suggests the initial formation of a partially stable bimolecular duplex construct, and the latter peak represents the GQ. The expected molecular mass of the GQ formed from the 8G construct is 3.3 x10^4^ Da and the measured value is consistent with this expectation.

The GQ structure formed from ssDNA strands containing a 6MI monomer (8Gm) or a 6MI dimer (8Gd) showed a single major peak around ∼ 3.2 x10^4^ Da and ∼ 3.1 x10^4^ Da, respectively, which suggests the formation of stable GQ structures had gone to completion (Figure 3C and D). Interestingly the peak observed at ∼ 1.9 x10^4^ Da for the unlabeled construct is absent in both the 6MI monomer- and the 6MI dimer-labeled constructs. This is consistent with the observations made with the gel electrophoresis assay, where the middle band corresponding to the bimolecular intermediate is absent in the 6MI monomer- and dimer-labeled constructs. Further, we notice that there is a slight decrease in the molecular weight of the GQ formed from the 6MI labeled construct. This is also in agreement with observations from gel-band experiments where a slight ‘spreading’ of the top band was observed. We speculate that the peak around ∼ 3.1-3.2 x10^4^ Da might be either coming from the sum of the trimer and tetramer structures, in accord with the EMSA results or due to the difference in hydrodynamic properties of the structures formed.

All these observations show that the 6MI monomer- or dimer-labeled ssDNA strands can form stable GQ complexes with the expected size and homogeneity similar to their unlabeled GQ counterpart (8G). The fact that we can make GQ structures with 6MI probes opens the possibility of site-specifically positioning these probes at positions of interest and monitoring the conformational change occurring at specific positions where crucial GQ-small molecule and GQ-protein interactions occur.

### Absorbance spectra of 6MI labeled ssDNA strands show hyperchromicity upon GQ formation

EMSA and sedimentation velocity experiments showed that eight consecutive guanines per ssDNA strand permit the formation of a highly stable parallel G-quadruplex structure with ssDNA strands that have site-specifically incorporated 6MI probes. Absorbance spectra of the free ssDNA strands in deionized water were compared with the spectra of the corresponding GQ constructs in 10mM Tris, pH 7.5 buffer and 100mM NaCl containing solutions to monitor the absorbance changes and identify the wavelengths where maximum changes are observed upon GQ formation with base analogue probe-containing ssDNA strands (Figure 4A). The experiments were performed at 20°C to avoid any unstacking effect and the resulting hyperchromicity due to temperature changes. The results showed that the formation of GQs resulted in a slight increase in absorbance at 260 nm for all constructs. Variation in the initial intensity of individual constructs at the absorbance maximum (260 nm) is due to the difference in the number of canonical bases. In 8Gm, one guanine is replaced by a 6MI; in 8Gd, two guanines are replaced by two 6MIs. Hyperchromicity upon formation of GQ is expected as the formation of GQ requires unstacking, rotation, and reorientation of guanines to form the Hoogsteen-base pairing. This hyperchromicity at 260 nm suggests that base stacking is more efficient in the ssDNA strands than in the final GQ structures.

**Figure 4.**
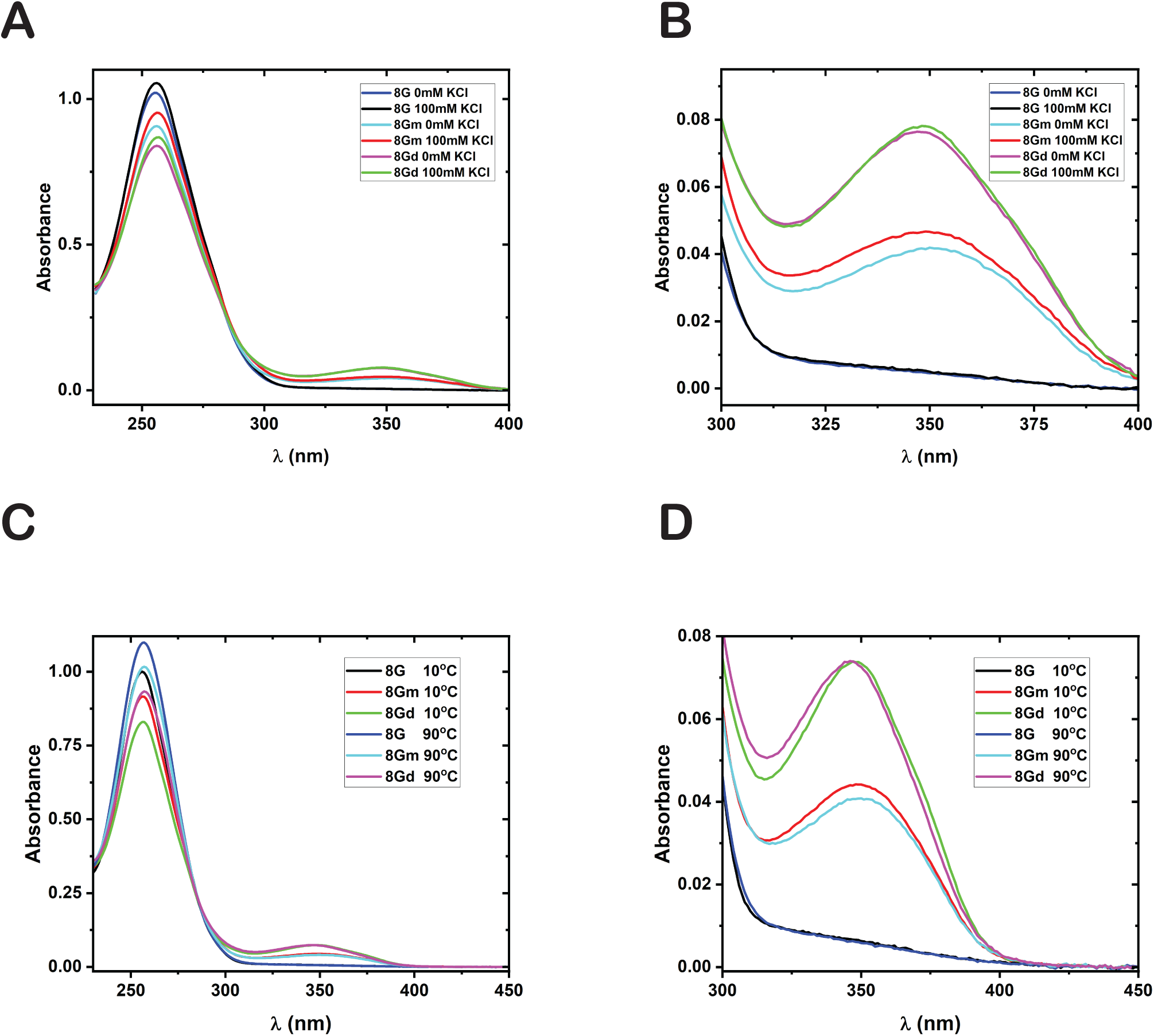
UV absorption spectra of unlabeled and 6MI monomer or dimer labeled GQ constructs at various salt concentrations and temperatures. (A) Absorbance spectra of the canonical base region and 6MI region and (B) only the 6MI region in no salt and with 100mM KCl at 20°C. (C) Absorbance spectra displaying the canonical base region and 6MI region and (D) only the 6MI region in 100mM KCl at 10°C and 90°C. All experiments were performed in 10mM Tris, pH 7.5 buffer, and at the listed salt concentrations.

The 6MI incorporated strands displayed an absorption band in the 325 nm to 375 nm region, a region where canonical bases or proteins do not show any absorbance (Figure 4B). In this noncanonical region, for the 8Gm construct, GQ formation resulted in a detectable enhancement of absorbance near the absorbance maximum of 6MI, ∼350 nm (Figure 4B). This opens the possibility of following the conformational changes of individual G4 layers formed by 6MI probes using thermal denaturation and other absorbance methods. However, for the 8Gd construct, the observed enhancement of absorbance at ∼350 nm is minimal. The lack of hyperchromicity indicates that the stacking between 6MI dimers is very strong. We speculate that the 6MI dimers are rotating and reorienting pairwise rather than as individual bases with minimal or hardly any 6MI-6MI unstacking.

### Monitoring the thermal unfolding of individual G4 layers by following the absorbance changes

Absorbance spectra of GQs formed from 8G, 8Gm, and 8Gd constructs were collected from 230 nm to 450 nm at 10°C and 90°C to verify the possibility of monitoring the thermal denaturation of individual G4 layers (Figure 4C, D). At 260 nm, where canonical bases melt, hyperchromicity is observed for all three constructs due to the thermal unstacking of bases. For 6MI-labeled constructs, a hyperchromic effect near 350 nm was expected that would correspond to the unstacking of 6MI probes. However, for construct 8Gd, no significant absorbance change was observed in the 350 nm region. This is consistent with the observation made in Figure 4B, where for construct 8Gd no significant change is observed in the 350 nm region when GQ is formed from its component ssDNA strands, suggesting the G4 layers formed by the stacking of two 6MI tetrad layers might be more stable than the tetrad layers formed by layers containing guanine residues only. Interestingly, absorbance spectra of 8Gm, where a 6MI G4 layer is stacked between two guanine tetrad layers, showed hypochromicity at 350 nm with an increase in temperature (Figure 4D). This indicates that 6MI monomer probe in the tetrad layer of 8Gm GQ complex is more unstacked than in a fully thermally denatured 8Gm construct. Therefore, we decided to monitor the change in absorbance of the 8Gm GQ at 260 nm and at 350 nm to elucidate the difference in structure and stability at the global and the local conformational levels.

Thermal melting experiments were performed on GQs formed from all constructs at 260 nm and 350 nm (Figures 5 and S1). At 260nm, all constructs showed a linear increase in absorbance with an increase in temperature with no defined baselines, making melting point determination impossible (Figure S1). We hypothesize that stacking eight continuous guanines (or guanines and 6MIs) makes the GQ structure extraordinarily stable and cannot be unfolded completely even at 90°C. Analysis of the thermal melting data at 350 nm, where the 6MI absorbs, showed different patterns for 8Gm and 8Gd constructs (Figure 5). The observed decrease in absorbance for 8Gd construct is not significant enough to monitor, whereas the 8Gm construct showed a significant, detectable decrease. This difference agrees with the absorbance spectra of the GQs from these two constructs taken at 10°C and 90°C (Figure 4D). We attribute this difference in melting pattern to the fact that the stacking of 6MI-6MI dimer is highly efficient compared with 6MI-G stacking. So even at elevated temperatures, the 6MI dimer must have remained stacked and thus showed little or no absorbance change (Figure 5C). This points out the limitation of using 6MI dimers as a probe in experiments aimed at monitoring the stability of GQ and other canonical and noncanonical nucleic acid structures. For 8Gm, the detectable decrease in absorbance was enough to determine the melting temperature of the fluorescently labeled 6MI tetrad layer (∼51°C, Figure 5A, Table 2). A comparison of melting profiles of 8Gm at 260 nm and 350 nm showed that the individual 6MI tetrad layer melted before the global melting of GQ structure. This suggests that the stability of GQ structures is the result of contribution from each individual G4 layer. The detection of the thermal denaturation of 6MI tetrad layer without global denaturation overlap revealed that the introduction of site-specific fluorescent base analogues that can form a G4 layer could be used to monitor the conformation and stability of individual G4 layers in a GQ complex.

**Figure 5.**
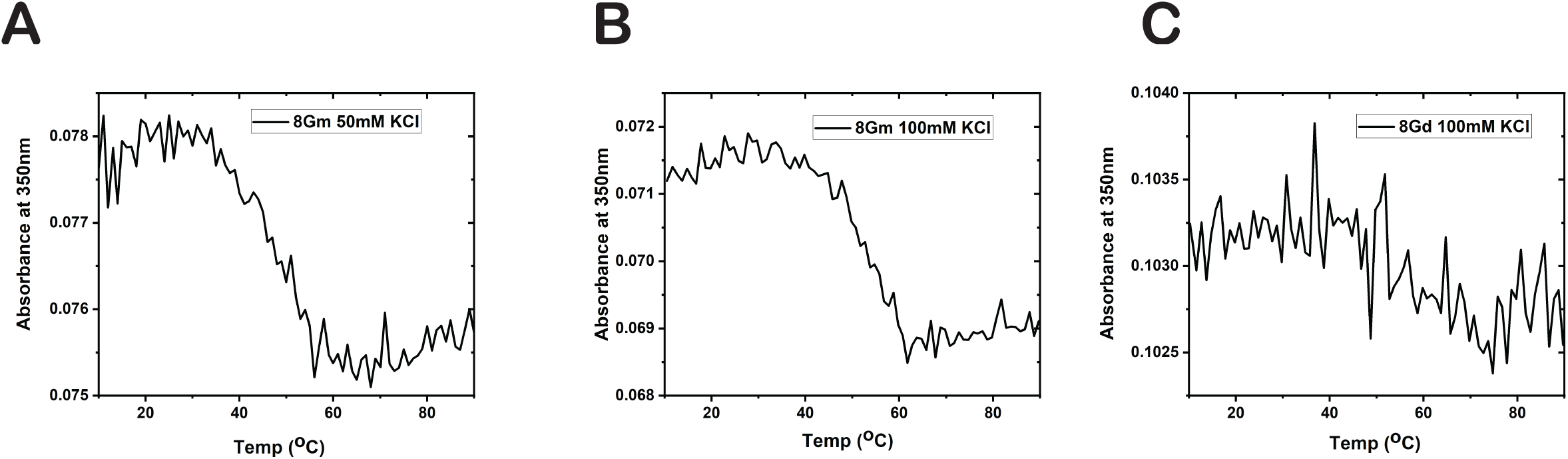
Melting curves of fluorescent G4 layers monitored by change in absorbance at the 6MI absorbance maximum, 350 nm. (A) UV thermal denaturation curves of the 6MI monomer labeled 8Gm GQ construct in 50mM KCl, 10mM Tris pH 7.5 buffer at 350 nm. (B) UV thermal denaturation curves of the 6MI monomer labeled 8Gm GQ construct in 100mM KCl, 10mM Tris pH 7.5 buffer at 350 nm. (C) UV thermal denaturation curves of the 6MI dimer labeled 8Gd GQ construct in 100mM KCl, 10mM Tris pH 7.5 buffer at 350 nm.

### 6MI labeled ssDNA construct show an enhancement in fluorescence intensity upon GQ formation

Previous studies showed that stacking of 6MI with canonical bases results in the quenching of fluorescence intensity of 6MI monomers or dimer probes[24,25]. The few exceptions to this general observation were observed with some specific sequences where thymine is the flanking base [26]. Further, it is known that 6MI-6MI stacking quenches the fluorescence more strongly than 6MI-Guanine stacking. The fluorescence intensity measurements of the 6MI labeled ssDNA constructs shown in Figure 6 demonstrate that the monomer-labeled unannealed 8Gm ssDNA has a higher fluorescence intensity than the dimer labeled 8Gd ssDNA construct, thus confirming the previous observations. Fluorescence spectra of annealed GQ structures were recorded in 10mM Tris buffer with varying salt concentrations to monitor the fluorescence intensity changes upon the formation of GQ structures. At 100mM KCl, where a stable GQ structure was expected, 8Gm constructs showed an ∼ 12-fold enhancement of fluorescence intensity, whereas the 8Gd constructs showed an ∼ 6-fold enhancement compared to the ssDNA constructs with a blue shift in the emission peak (Figure 6A, B). Further, it was noted that for the 8Gm construct, the fluorescence intensity showed an increase with an increase in salt concentration, however for the 8Gd construct the fluorescence intensity observed at 50mM KCl concentration is higher than that at 100mM KCl.

**Figure 6.**
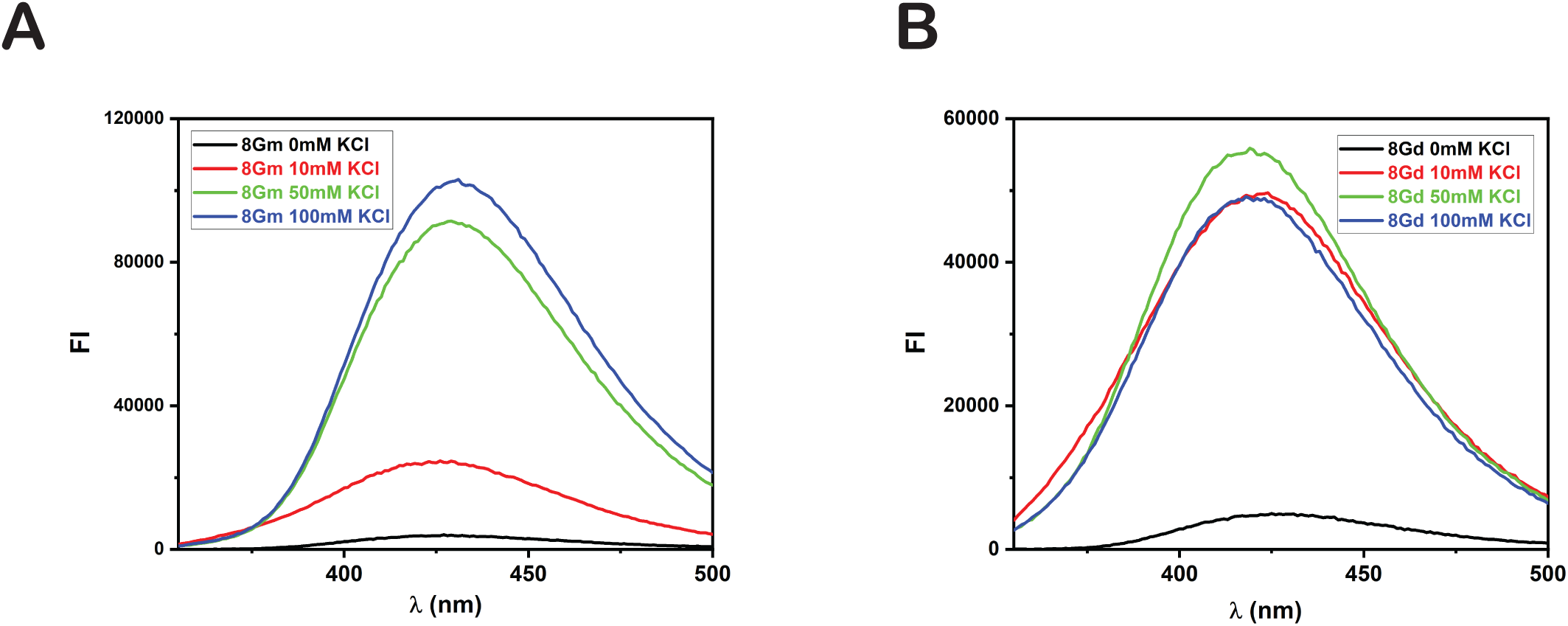
Fluorescence spectra of GQ constructs with either 6-MI monomer, 8Gm (A) or dimer, 8Gd (B) bases at defined positions at various salt concentrations. 8Gm constructs showed an enhancement of fluorescent intensity with an increase in salt concentration, whereas for 8Gd construct maximum fluorescence intensity was observed at 50mM KCl.

The formation of GQs is associated with stacking bases of the G4 layers and therefore expected to cause a decrease in fluorescence intensity compared with the unfolded structure. However, as discussed in the Observed Hyperchromicity in Absorbance with GQ Formation section above, rotation and reorientation of bases to form the G4 layers will result in unstacking individual bases from their neighbors. This partial unstacking of neighboring bases will diminish the quenching of 6MI fluorescence intensity, causing a fluorescence enhancement in GQ formation. The fluorescence and absorbance results show that the stacking of bases is more efficient in ssDNA than in GQ and that the extra stability of the GQ structure mainly comes from the cation coordination of the structure. This observation further emphasizes the importance of the cation involved in the structure and stability of GQs. Additionally, it is known that the dipole of guanines will change upon GQ formation, and the observed blue shift in the emission peak is attributed to the change in the dipole of individual 6MI residues with the formation of quadruplexes. The enhancement of fluorescence intensity upon GQ formation showed that the site-specific incorporation of fluorescent probes could monitor the local conformations of G4 layers. This method can also be used with absorbance methods to monitor the conformational properties of individual G4 layers in GQ structures.

### Monitoring the thermal unfolding of individual G4 layers by fluorescent intensity changes

After successfully incorporating a fluorescent G4 layer into the GQ and characterizing its fluorescence intensity fluctuations under different conditions, our next aim was to develop a method to monitor the conformation and stability of individual G4 layers. The synthetic sequence used in this study is known to form a highly stable GQ, and a complete denaturation of the GQ structure was not observed even at 90°C in absorbance measurement (Fig S1). For the 8Gm constructs in 100mM KCl, the absorbance change observed at 350 nm, the 6MI probe’s absorbance maximum, was significant enough to monitor the changes. However, at lower salt concentrations, the observed decrease in absorbance was not substantial for accurately determining the melting point. Fluorescence intensity measurements showed a significant increase with GQ formation, so we decided to monitor the fluorescence intensity variations of the GQ structure at different salt concentrations. We hypothesize that since the stability of GQs varies with salt concentration, the thermal denaturation properties of the fluorescent G4 layer also might show a corresponding variation.

Thermal denaturation experiments were performed at three different salt concentrations for both 8Gm and 8Gd constructs (Figure 7). For 8Gm constructs, the increase in temperature resulted in a decrease in fluorescence intensity at all salt concentrations (Figure 7A). This is in line with the results observed previously, as the formation of GQ showed an enhancement in fluorescence intensity of the probe upon GQ formation. Therefore, as the denaturation occurs, the highly fluorescent GQ structure collapses and forms the less fluorescent single strand structure. This agrees with the previously made observation that GQ formation from ssDNA construct caused an ∼ 12-fold enhancement in fluorescence intensity. Further, the results showed that an increase in salt concentration raised the melting point of the GQ constructs (Table 2). These observations indicate that a fluorescent G4 layer formed by properly incorporated 6MI monomers can be used to monitor the local conformation of individual G4 layers in a GQ structure. However, the results observed for the 8Gd constructs showed an increase in fluorescence intensity with an increase in temperature. Even at 90°C, the fluorescence intensity did not reach a plateau suggesting unstacking of the bases from neighboring bases is not yet complete (Figure 7B). It is known from previous studies that 6MI dimer probes exhibit large self-stacking and form highly stable dimers [24]. For example, helicases and polymerases that usually unwind canonical bases and other fluorescent base analogues, including 6MI monomers, cannot act on 6MI dimer probe [24]. In the present case, an increase in temperature will cause the unstacking of the highly stacked 6MI-6MI dimers, decreasing the self-quenching and increasing fluorescence intensity. This decrease in self-quenching and the resulting fluorescence enhancement overshadow the reduction in fluorescence intensity occurring during the collapse of the GQ structure.

**Figure 7.**
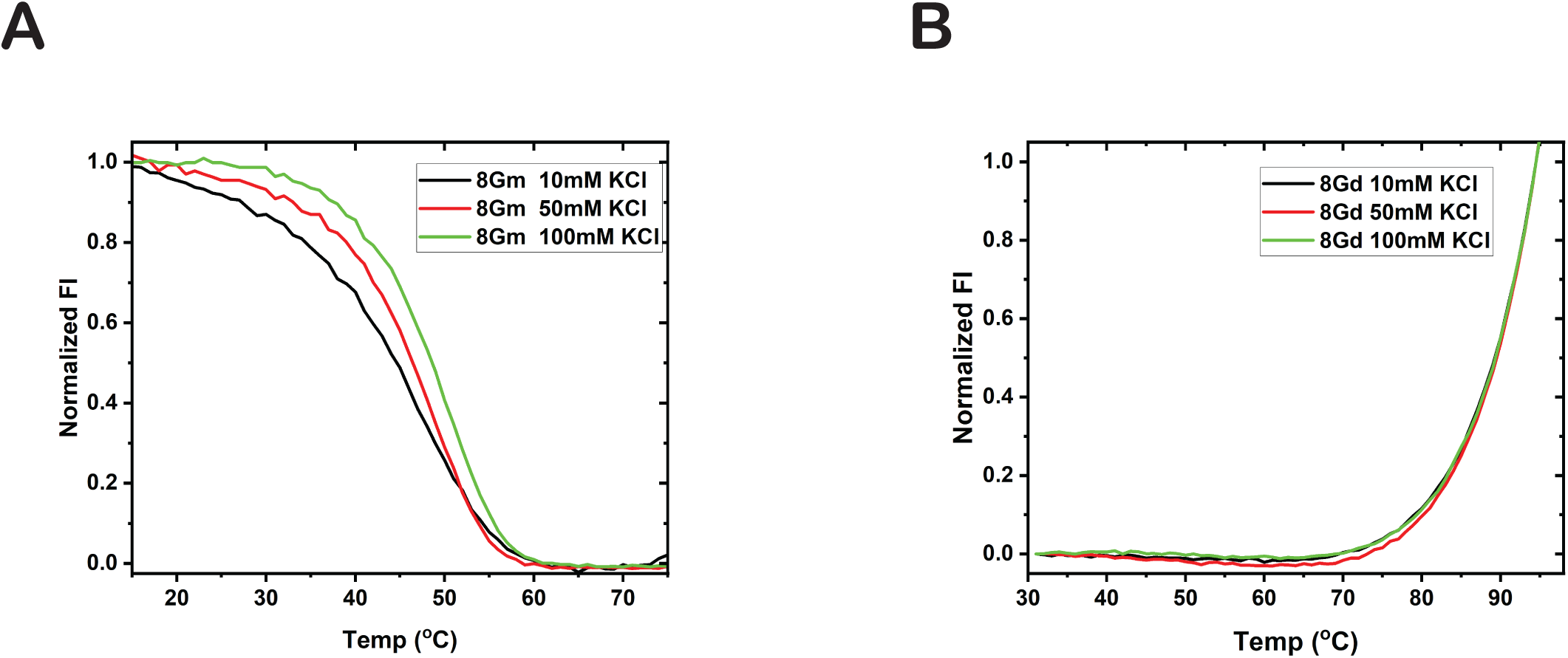
Melting curves of fluorescent G4 layers. (A) Fluorescence melting curves at the emission maximum wavelength 430nm (excitation at 350nm) for 8Gm (A) and 8Gd (B) at different salt concentrations.

### Fluorescence quenching experiments showed the G4 layers in GQ are more accessible to solvent than in the ssDNA construct

The quenching of fluorescence of base analogues in DNA by acrylamide depends on the solvent accessibility of fluorophores. Previously, we used this approach to acquire specifics of structural and dynamic aspects of DNA breathing near ss/ds junctions[29,41,46]. A higher quenching coefficient indicates more solvent accessibility, and we can monitor the differences in solvent accessibility of the probe with variation in structural stability and conformational differences by this method. Acrylamide fluorescence quenching of 6MI residues in GQ was performed on 8Gm constructs to provide additional insight into the structure of the fluorescent G4 layer. The results were fitted to obtain the Stern-Volmer (S-V) quenching constants, and the parameters obtained are summarized in Table 3.

GQ structures from the 8Gm construct were formed at 50mM or 100mM KCl in Tris buffer to perform quenching experiments. The addition of acrylamide at 20°C resulted in fluorescence quenching at both salt concentrations indicating solvents can access the G4 layer of the GQ structure (Figure 8A). The Stern-Volmer plot is linear, suggesting the occurrence of only one type of quenching type, either dynamic or static (Figure 8B). To differentiate between static and dynamic quenching, experiments were performed at 50°C. An increase in temperature increased the quenching constant, confirming collisional quenching was occurring (Figure 8 C, D, Table 3). The larger quenching constant (Ksv) observed for the construct at 50mM KCl compared to that at 100mM KCl confirms that the increase in salt concentration stabilizes the GQ structure, and the solvent accessibility decreases as the stability of the structure increases. This experiment proved that the internal G4 layer is accessible to solvents, and the degree of accessibility varies with the stability of GQs. In the future, this approach can be used to determine the relative accessibility of the GQ layers to various (‘intercalating’?) small molecules that can be used as potential drugs for treating GQ-related diseases focused towards internal G4 layers and GQ structures as a whole.

**Figure 8.**
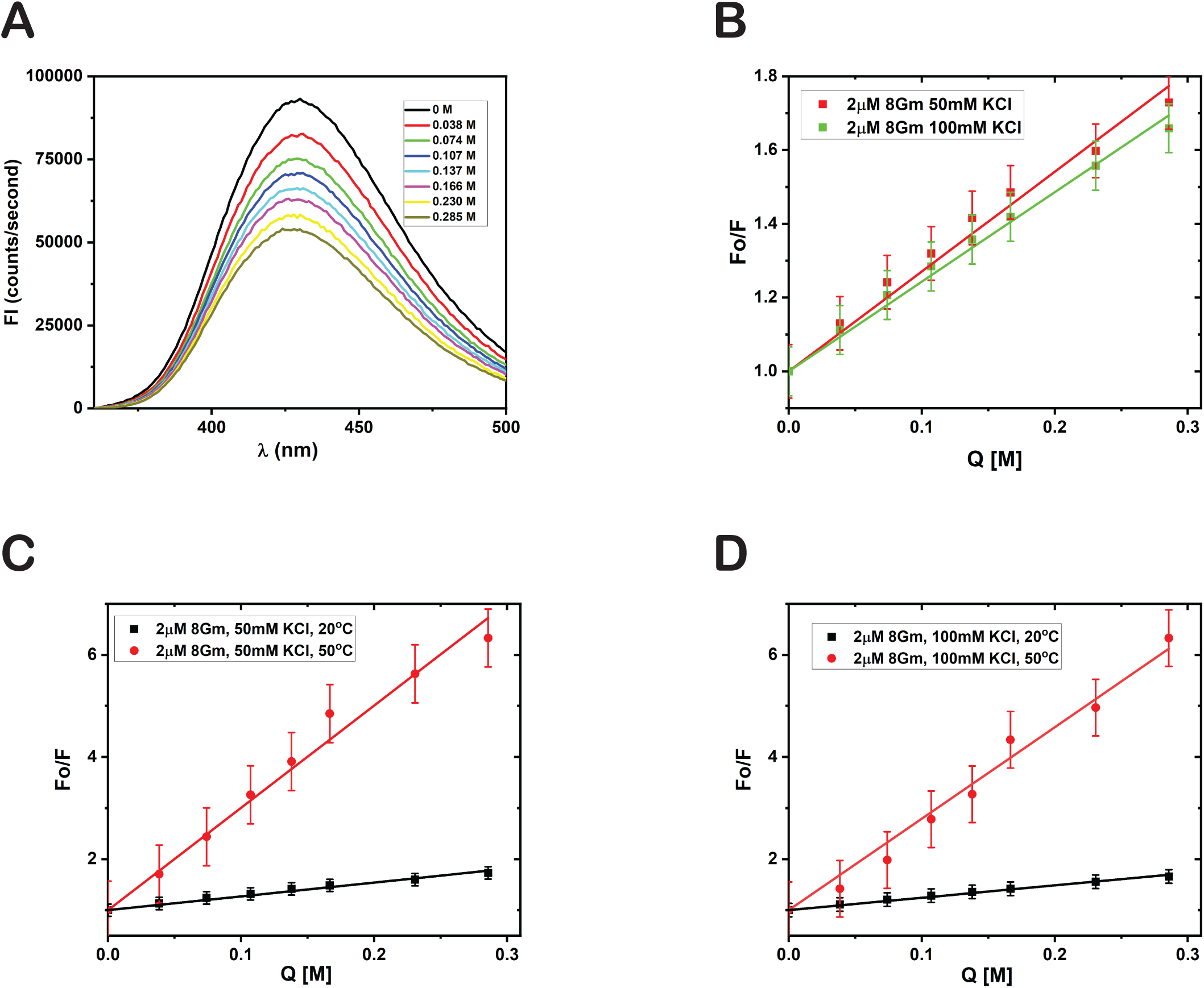
Acrylamide quenching of 6MI probes in GQ constructs. (A) Representative fluorescence spectra of 8Gm constructs in 100mM KCl, 10mM Tris pH 7.5 buffer with increasing acrylamide concentration. (B) Stern-Volmer plots for 8Gm GQ with 50mM and 100mM KCl concentrations titrated with monomeric acrylamide as the quenching agent at 20°C. Comparison of quenching at 20°C and 50°C at 50mM (C) and 100mM (D) KCl concentrations.

### 6MI dimer-labeled ssDNA construct show characteristic low energy CD signal upon GQ formation

Earlier studies with CD-active fluorescent base analogues, especially 2AP and 6MI, showed that CD spectroscopy of dimer probes could provide local structural information that is not obtainable with monomer probes [24, 29]. Experiments were performed with unannealed 8G, 8Gm, and 8Gd constructs in 0mM KCl and with annealed constructs in 100mM KCl to monitor the CD signal changes when GQs are formed from ssDNA (Figure 9A). All constructs showed the characteristic parallel GQ peak (∼ 265 nm) and trough (∼235 nm) in the canonical base region of 230 nm-300 nm. The variation in the intensity of the peaks is attributed to the difference in the number of canonical bases. As seen in Figure 9A, 8G constructs with 26 canonical bases in the monomer strand showed the highest intensity as the unannealed ssDNA and GQ constructs, whereas 8Gd with 24 canonical bases in the monomer strand showed the least. The formation of GQs from ssDNA constructs resulted in the increase of CD signal for all constructs, and this further confirms that the presence of 6MI bases does not hinder the formation of GQs. Analysis of the low energy CD region, 300-450 nm, showed detectable signals only for the 8Gd constructs (Figure 9B). The ssDNA construct showed a trough at ∼ 335nm, and this trough disappeared as the GQ was formed. In addition, the formation of the GQ showed the appearance of a characteristic peak near 375 nm. However, the CD spectrum of 8Gm ssDNA and the corresponding GQ construct did not show any detectable CD signal in the low energy 300 to 400 region. This is because, in the case of 6MI monomers, the exciton coupling is between the 6MI and the neighboring canonical bases, and the resulting CD signal is not with detectable intensity. Therefore, it is impossible to track the relative local conformation of the single G4 layer in the GQ using 6MI monomer probes using CD spectroscopy. Conversely, for the 8Gd construct, the ssDNA and the corresponding GQ showed a detectable CD signal in the low-energy CD region characteristic of exciton coupled 6MI dimer probes. We previously showed that under favorable conditions, the exciton coupling of 6MI dimers would result in a CD spectrum that can provide specific information about the relative conformation of the individual bases in the dimers without the overlap of signals from the neighboring bases [24, 25].

**Figure 9.**
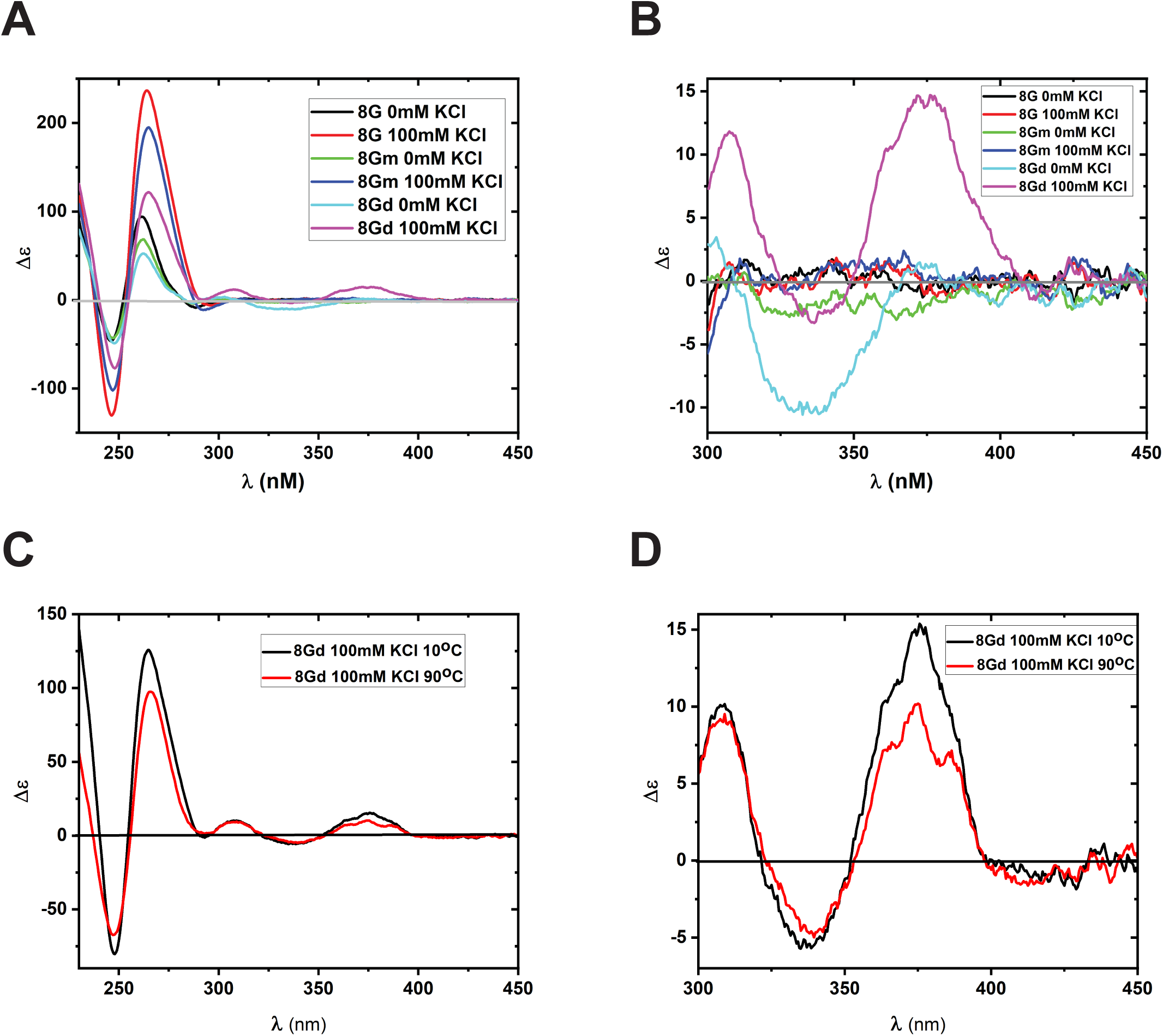
CD spectra of GQ constructs with either 6-MI monomer and dimer bases at defined positions. (A) CD spectra of 8G, 8Gm, and 8Gd constructs in the single strand (unannealed) and after forming GQ (annealed) state. (B) CD spectra showing the signal only from the 6MI probes. (C) CD spectra of 8Gd constructs at 10°C and 90°C showing both canonical and noncanonical regions and (D) showing the signal only from the 6MI probes (noncanonical region).

To gather more data about the relative structure and orientation of adjacent G4 layers of a GQ, the thermal unfolding of GQs of 8Gd constructs were explored (Figure 9C, D). In the canonical region increase in temperature resulted in a decrease in intensity for the peak (265nm) and trough (245nm) (Figure 9C). In the noncanonical region, the peak at 375nm decreased; however, the trough at 335nm remained more or less unchanged (Figure 9D). Melting profiles of the GQ constructs were then monitored at 265nm for the canonical region (Figure S2A) and 375nm for the noncanonical region (Figure S2B). The data and the resulting plot obtained are not conclusive enough to precisely predict the melting point of either the global or local conformation of the GQ. This is in accordance with what was observed in absorbance and fluorescence thermal-unfolding experiments. The high stability of the sequence used in this study might be hindering the thermal denaturation of 8Gd constructs. Our ongoing investigations with human telomeric GQ sequence are exploring different methods to extract details of thermal stability, and the results look promising as the GQ structures contain only three G4 layers.

## CONCLUSION

G-quadruplexes are biologically relevant noncanonical secondary nucleic acid structures that are involved in many cellular processes such as DNA replication, transcription, recombination, epigenetics and telomeric stability. Studies on the structure, stability, and interaction with small molecules of GQs were always confined to the global conformation. In this study, we utilized several complementary spectroscopic methods such as absorbance, thermal denaturation, fluorescence intensity, fluorescence quenching, and CD experiments to explore the local conformations of individual G4 layers of a GQ structure. The results showed that it is possible to form an internally fluorescent GQ by incorporating site-specifically placed 6MI probes. The formation of GQ is associated with an increase in absorbance and fluorescence intensity which can be used to monitor the formation of GQ in vitro for experimental studies. To form G4 layers, individual bases should unstack from their neighboring bases in the ssDNA and undergo rotation and reorientation to form the Hoogsteen base pairing. This structural change required for the formation of GQ causes an increase in absorbance and fluorescence upon GQ formation. Of course, these experiments are performed on a synthetic DNA strand, and currently, we are performing experiments on human telomeric sequences and several other potentially GQ-forming sequences to see if different results are obtained with *in ‘vivo’* systems. A detailed understanding of the structure of individual G4 layers and their interactions with small molecules should eventually help in the development of new therapeutic drugs and thus suggest new methods to treat many telomere and GQ-related diseases such as Bloom, Werner, and other cancer-type syndromes.

## ASSOCIATED CONTENT

The Supporting information is available at free of charge at Supplementary Figures S1 and S2

## FUNDING

Monmouth University Creativity and Research Grant to Dr. Jose.

## ACKNOWLEDGEMENTS

We thank Dr. Peter H von Hippel for many useful discussions of this work and for permitting us to use the Analytical Ultracentrifuge at University of Oregon. We thank Steven Weitzel for setting up and running the AUC. We are also very grateful to Dr. Walter Baase for careful readings of the manuscript. We thank Dr. Jonathan Ouellet for many helpful discussions throughout the course of this project.

## Notes

### Competing Interest Statement

The authors have declared no competing interest.

### Summary of Updates

Abstract, Introduction,Results and Discussions updated

