## Supplementary material for "A Spectroscopic Approach to Unravel the Local Conformations of G-quadruplex Using CD-active Fluorescent Base Analogues": Supplemetary Figures1 and 2

**SUPPORTING INFORMATION**

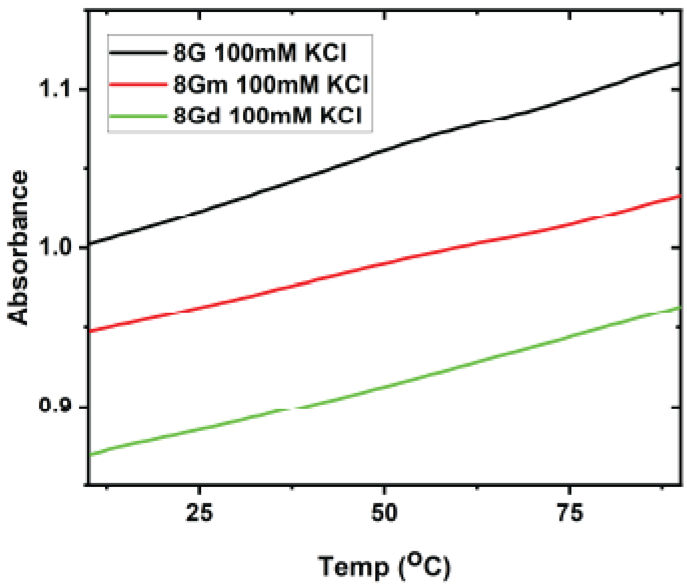


**Figure S1. Thermal denaturation curves monitored by the change in absorbance at 260 nm.** (A) UV thermal denaturation curves of the unlabeled 8G, 6MI monomer labeled 8Gm and 6MI dimer labeled 8Gd GQ constructs in 100mM KCl, 10mM Tris pH 7.5 buffer at 260 nm.

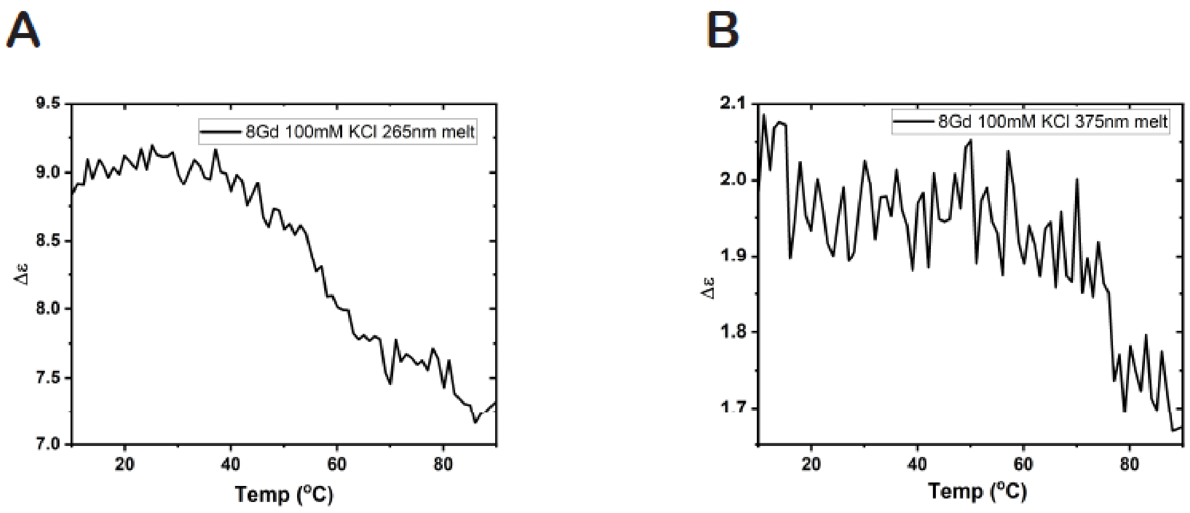


**Figure S2. Thermal denaturation curves monitored by CD changes of 8Gd constructs at 265 nm and 375 nm.** (A) CD thermal denaturation curves of 6MI dimer labeled 8Gd GQ constructs in 100mM KCl, 10mM Tris pH 7.5 buffer at 260 nm and (B) at 375 nm.
